# Draft genome assemblies using sequencing reads from Oxford Nanopore Technology and Illumina platforms for four species of North American killifish from the *Fundulus* genus

**DOI:** 10.1101/686246

**Authors:** Lisa K. Johnson, Ruta Sahasrabudhe, Tony Gill, Jennifer Roach, Lutz Froenicke, C. Titus Brown, Andrew Whitehead

## Abstract

Draft *de novo* reference genome assemblies were obtained from four North American killifish species (*Fundulus xenicus, Fundulus catenatus, Fundulus nottii*, and *Fundulus olivaceus*) using sequence reads from Illumina and Oxford Nanopore Technologies’ PromethION platforms. For each species, the PromethION platform was used to generate 30-45x sequence coverage, and the Illumina platform was used to generate 50-160x sequence coverage. Contig N50 values ranged from 0.4 Mb to 2.7 Mb, and BUSCO scores were consistently above 90% complete using the Eukaryota database. Draft assemblies and raw sequencing data are available for public use. We encourage use and re-use of these data for assembly benchmarking and external analyses.

## Background

Sequencing and assembling large eukaryotic genomes is challenging [1–3]. Accuracy of downstream analyses, such as variant calling and measuring gene expression, depends heavily on a high-quality reference genome [4]. Fortunately, the cost of generating whole genome sequence data is dropping, making it easier for individual labs rather than large consortiums to generate assemblies for organisms without reference genomes [3,5,6]. Single-molecule long read nucleic acid sequencing technology from Oxford Nanopore Technologies (ONT), which has been commercially available since 2014 [7], has been shown to improve the contiguity of reference assemblies [8] and reveal “dark regions” that were previously camouflaging genes [9]. The lengths of the sequencing reads generated using this technology are limited only by the size of the fragments in the extracted DNA sample [10]. The promise of more complete reference assemblies is especially important for the accuracy of comparative evolutionary genomics studies, as assembly fragments lead to errors in downstream synteny analyses [11], as well as SNP calling and identification of transcript features (splice junctions and exons) for quantification.

Despite high error rates ∼5% [12] relative to Illumina short reads ∼0.25% [13] and the relatively recent availability of ONT data, there have been a flurry of studies using this sequencing technology. Small genomes from bacteria and viruses appear to be ideal for sequencing on the ONT MinION platform [12]. The portable nature of the technology makes it appealing as a resource for teaching [14,15], working in remote locations [16–18] and for investigating viral outbreak public health emergencies [19–21]. However, despite the demonstrated ability to achieve yields >6.5 Gb per flow cell [22], the MinION platform can be prohibitively expensive for sequencing larger eukaryotic genomes. For example, 39 flow cells yielded 91.2 Gb of sequence data (∼30x coverage) of the human genome [23]. Sequencing of the wild tomato species, *Solanum pennellii* across thirty-one flow cells yielded 110.96 Gb (∼100x coverage) with some flow cells yielding >5Gb [24]. By contrast, following the 2018 beta release of the ONT PromethION platform, which has a higher density of nanopore channels, five flow cells were used to yield >250 Gb (∼80x coverage) of the human genome [25]. PromethION data combined with Hi-C long-range mapping data from human samples produced a genome genome assembly with a scaffold N50 of 56.4 Mbp [26].

The combination of long read sequencing data from ONT MinION and short read sequencing data from Illumina has been used to improve the quality of reference genomes [27–30]. In one approach, short read assembly scaffolds have been improved with the addition of long reads. The Murray cod genome (640-669 Mb in size) was improved by combining low coverage 804 Mb of long reads ONT data from just one MinION flow cell with 70.6 Gb of Illumina data from both HiSeq and MiSeq; the assembly scaffold N50 increased from 33,442 bp (Illumina only) to 52,687 bp with ONT and Illumina combined [31]. The clownfish genome (791 to 794 Mb in size) was improved by including 8.95 Gb of ONT MinION reads; the scaffold N50 increased from 21,802 bp (Illumina only) to 401,715 bp with ONT and Illumina combined [27]. Recently, a new approach is available with racon [32] and/or pilon [33] consensus building tools, which uses Illumina data to “polish” contigs from ONT-only assemblies. Polishing corrects single nucleotide base differences, fills gaps, and identifies local mis-assemblies [33]. This approach has been shown to improve the BUSCO score from <1% with the ONT assembly alone to >95% complete after polishing with Illumina reads [28].

In this study, we explored whether the ONT PromethION sequencing technology could be appropriate for generating initial draft reference genomes for four species of North American killifish belonging to the *Fundulus* genus. *Fundulus* is a comparative evolutionary model system for studying repeated genomic divergence between marine and freshwater species. *Fundulus* killifish have a cosmopolitan geographic distribution across North America. These small cyprinodontiform fish have evolved to occupy a wide range of osmotic niches, including marine, estuarine, and freshwater [34]. Estuarine and coastal *Fundulus* are euryhaline, insofar as they can adjust their physiologies to tolerate a very wide range of salinities. In contrast, freshwater species are stenohaline: they tolerate a much narrower range of salinities [34,35]. Freshwater clades are derived from marine clades, and radiation into freshwater has occurred multiple times independently within the genus. This makes *Fundulus* unusual, because most large clades of fishes are either exclusively marine or exclusively freshwater. Therefore, species of closely-related killifish in the *Fundulus* genus serve as a unique comparative model system for understanding the genomic mechanisms that contribute to evolutionary divergence and convergence of osmoregulatory processes, which is important for understanding how species will evolve to face fluctuating salinities in future climate change scenarios [36]. The Atlantic killifish, *Fundulus heteroclitus* has been a well-described physiological model organism for investigating the functional basis of, and evolution of, physiological resilience to temperature, salinity, hypoxia, and environmental pollution [34,37–39], with an available genome from the Atlantic killifish, *Fundulus heteroclitus* [40]. However, we do not currently have any genomes from other *Fundulus* killifish, particularly from those occupying freshwater habitats.

Here, we report the collection of whole genome sequencing data using both ONT PromethION and Illumina platforms from four killifish species without previously-existing sequencing data (Figure 1): *Fundulus xenicus* (formerly *Adinia xenica*) [41], *Fundulus catenatus, Fundulus nottii*, and *Fundulus olivaceus. F. xenicus* is euryhaline and occupies coastal and estuarine habitats, while the other species (*F. catenatus, F. nottii, F. olivaceus*) are stenohaline and occupy freshwater habitats.

**Figure 1.**
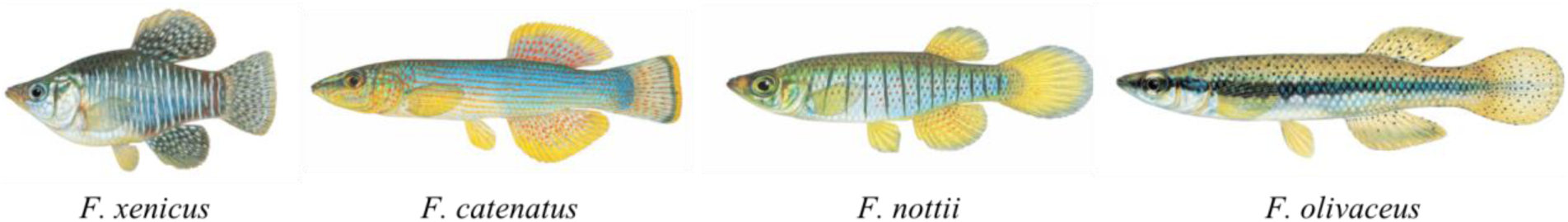
Four *Fundulus* killifish (left to right): the marine diamond killifish *Fundulus xenicus*; the northern studfish, *Fundulus catenatus* (south central United States); the freshwater bayou topminnow, *Fundulus nottii*; and the freshwater blackspotted topminnow, *Fundulus olivaceus*. (drawings used with permission from the artist, Joseph R. Tomelleri).

## Methods and Results

Live field-caught individuals of each fish species were shipped to UC Davis and kept at their native salinities in an animal holding facility, maintained according to University of California IACUC standards. *F. catenatus* and *F. olivaceus* were collected from the Gasconade River, MO (latitude/longitude coordinates 37.879/-91.795 and 37.19/-92.56, respectively), *F. nottii* was collected from Walls Creek, MS (31.154433/-89.245381), and *F. xenicus* was collected from Graveline Bayou, MS (30.368756/-88.719329). High molecular weight (hmw) DNA was extracted from fresh tissue for *F. nottii* and *F. xenicus*, and from frozen tissue for *F. catenatus* and *F. olivaceus*. For *F. catenatus* and *F. olivaceus*, tissues were dissected and frozen in liquid nitrogen and stored at -80 °C until samples were prepared for hmw DNA extraction. With the exception of *F. olivaceus*, each assembly consisted of sequencing one tissue sample from one individual. For *F. olivaceus*, Illumina data were collected from DNA extracted from one individual while the ONT PromethION data were collected from another individual (frozen tissue).

### DNA extractions

Whole fish heads were used for hmw DNA extractions. Agilent’s Genomic DNA Isolation kit (Catalog #200600) was used to extract DNA from fresh tissues from *F. xenicus* and *F. nottii*. For *F. catenatus* and *F. olivaceus*, both the ultra-long read sequencing protocol from [42] (which included Tissue lysis buffer with Tris, NaCl, EDTA, SDS and Proteinase K followed by phenol:chloroform extraction), as well as the Qiagen “DNA purification from tissue using the Gentra puregene Tissue Kit” (p. 39) were used, and were found to be similar to the Agilent kit. Precipitated DNA was difficult to re-dissolve; therefore, additional phenol:chloroform cleanup steps were required after extractions. We found that adding urea to the lysis buffer helped the precipitated DNA pellet to be less fragile and go into solution easier [43]. Prior to library preparation, hmw DNA from *F. nottii* and *F. olivaceus* (PromethION) was sheared to 50 kb in an effort to improve the ligation enzyme efficiency, resulting in fragments in the 50-70 kb range. Field inversion gels were used to visualize hmw DNA (Figure 2).

**Figure 2.**
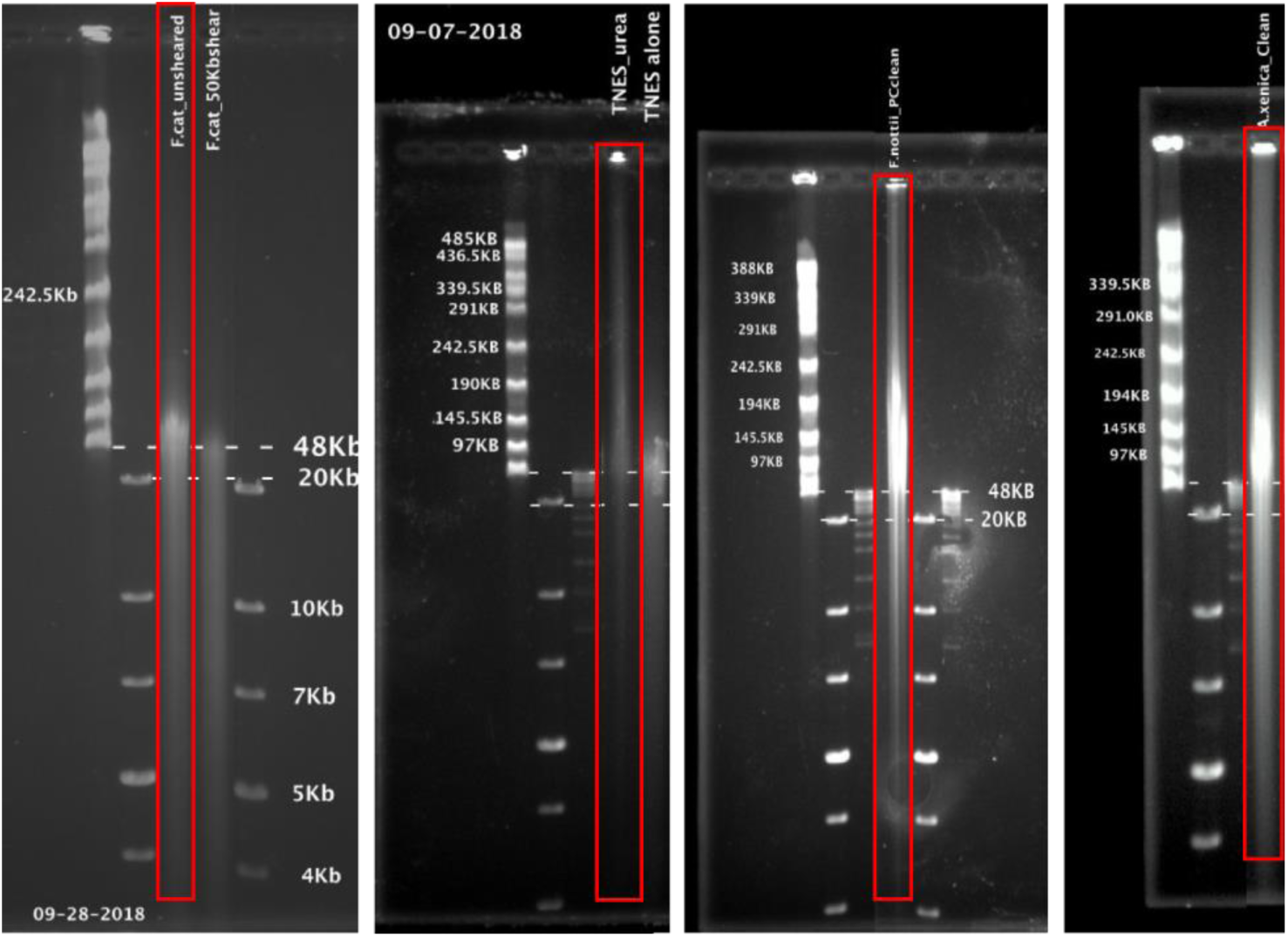
Field inversion gels with red boxes showing samples sequenced (in order from left to right: *F. catenatus* (sheared vs. unsheared), *F. olivaceus, F. nottii, F. xenicus*). DNA was extracted from fresh tissues for *F. xenicus* and *F. nottii*, and from frozen tissue for *F. catenatus* and *F. olivaceus.*

### ONT sequencing

Libraries for ONT PromethION sequencing were prepared using the ligation sequencing kit (SQK-LSK109) following the manufacturer’s instructions. ONT PromethION sequencing data were collected from all four species on an alpha-beta instrument through the early release program at the University of California, Davis DNA Technologies Core facility (Davis, CA USA). One species was sequenced per flow cell. Base-calling was done onboard the PromethION instrument (Oxford Nanopore Technologies, UK). For the *F. xenicus* run, lambda phage (DNA CS) was spiked-in as a positive control.

### Illumina Sequencing

With the exception of *F. olivaceus*, each individual hmw DNA sample used for the ONT library was also used for Illumina library preparation using the Nextera Index Kit (FC-121-1012). For three species, Illumina data were multiplexed across two PE150 lanes on an Illumina HiSeq 4000 and demultiplexed by Novogene (Sacramento, CA USA). For *F. olivaceus*, PE150 Illumina NovaSeq reads from one flow cell (2 lanes) were graciously provided by the Texas A&M Agrilife Research Sequencing Facility (College Station, TX USA).

## Data Description

Whole genome sequencing data from individuals of four killifish species collected from ONT PromethION (Table 1) and Illumina (NovaSeq and HiSeq 4000) (Table 2) were deposited in the European Nucleotide Archive (ENA) under the study accession PRJEB29136. Deposited raw data are untrimmed and unfiltered. Reads corresponding to lambda phage were filtered from ONT PromethION data using the NanoLyse program from NanoPack (version 1.1.0; [44]). Porechop (version 0.2.3) was used to remove residual ONT adapters and NanoFilt (version 2.2.0; [44]) was used to filter reads with an average quality score >Q5. After filtering and adapter trimming, ONT data from the PromethION ranged from 30-45x coverage for each species. NanoPlot (version 1.10.0; [44]) was used for visualization of ONT read qualities.

**Table 1.** ONT data collected from each species. Coverage assumes the genome size of each species is 1.1 Gb, measured for *F. heteroclitus* (Reid et al. 2017). Untrimmed reads were deposited in the ENA under study PRJEB29136. Q>5 reads were used in subsequent assemblies.

| Species | Bases called (Gb) | Cov. (x) | Average read length | Reads N50 | Q>5 Bases called (Gb) | Q>5 Avg. read length | ONT signal Accession | ONT fastq Accession |
| --- | --- | --- | --- | --- | --- | --- | --- | --- |
| *F. xenicus* | 38.5 | 35 | 2,449 | 5,733; n = 1,373,426 | 36.42 | 2,699 | ERR3385273 | ERR3385269 |
| *F. nottii* | 33.4 | 30.4 | 6,480 | 12,995; n = 700,534 | 31.06 | 7,548 | ERR3385275 | ERR3385271 |
| *F. catenatus* | 40.3 | 36.6 | 1,699 | 3,439; n = 2,687,295 | 34.28 | 2,021 | ERR3385274 | ERR3385270 |
| *F. olivaceus* | 50.1 | 45.5 | 4,595 | 11,670; n = 987,921 | 45.97 | 5,365 | ERR3385276 | ERR3385272 |

**Table 2.** Illumina data collected were all paired-end PE 150 reads. Coverage assumes 1.1 Gb genome size measured for *F. heteroclitus [40]*.

| Species | Platform | Reads (M) | Coverage (x) | FASTQ Accessions |
| --- | --- | --- | --- | --- |
| *F. xenicus* | Illumina HiSeq | 327.5 | 89.3 | ERR3385278<br>ERR3385279 |
| *F. nottii* | Illumina HiSeq | 197 | 53.7 | ERR3385282<br>ERR3385283 |
| *F. catenatus* | Illumina HiSeq | 316.5 | 86.3 | ERR3385280<br>ERR3385281 |
| *F. olivaceus* | Illumina NovaSeq | 601.9 | 164 | ERR3385284<br>ERR3385285 |

Average quality scores for all Illumina data were consistently above Q30 (Figure 3A). Residual Nextera adapters and bases with low quality scores were removed from Illumina reads using Trimmomatic PE (version 0.38) with conservative parameters, which included removing bases from each read with a quality score below Q2 and required a minimum read length of 25 bases each [45].

**Figure 3.**
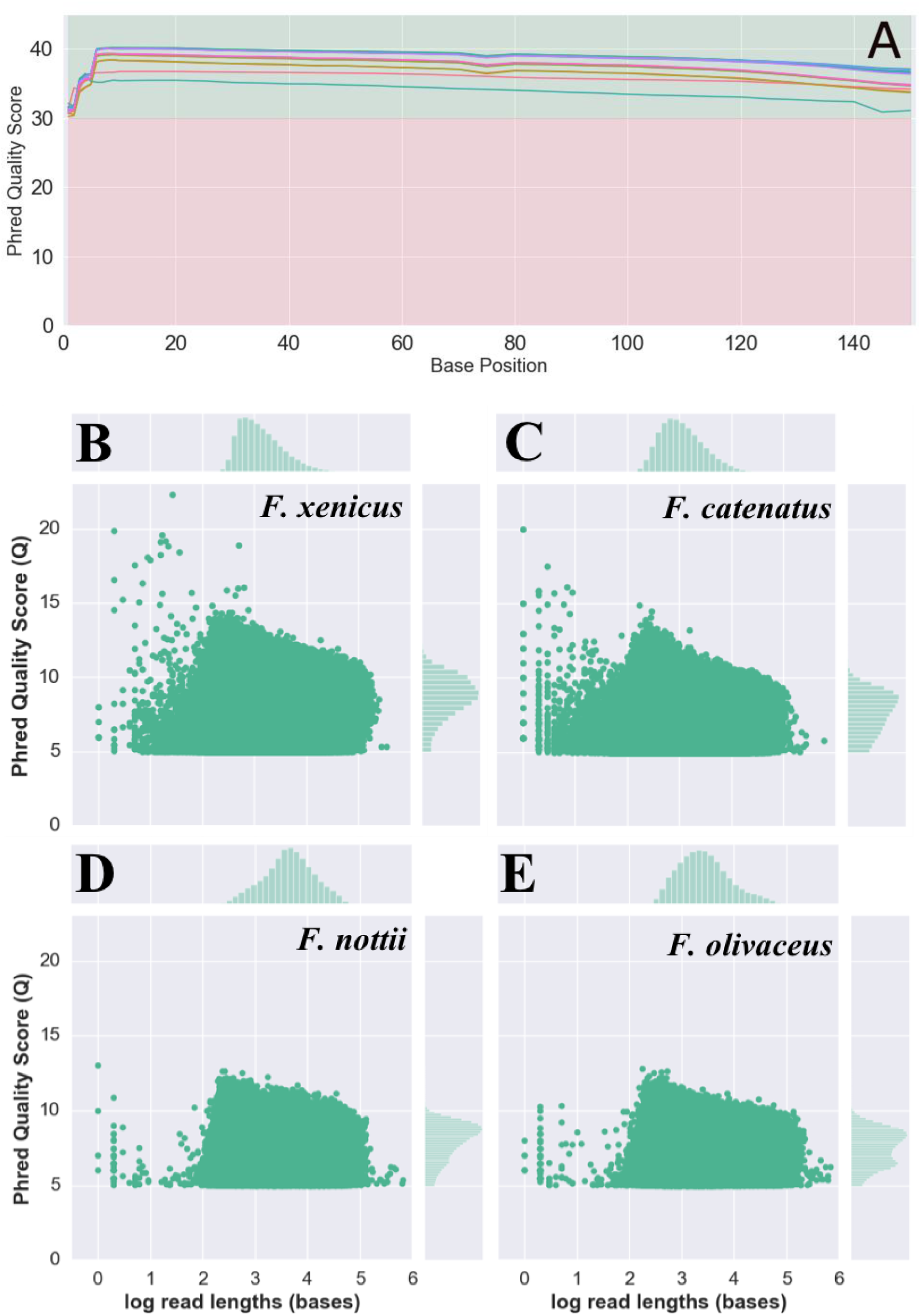
A) Quality score profiles for PE Illumina reads from *F. xenicus, F. catenatus, F. nottii* and *F. olivaceus.* For Illumina data, phred quality scores were consistently above Q30 across all reads. Average read quality scores (Q score) vs. read lengths for ONT PromethION from B) *F. xenicus*, C) *F. catenatus*, D) *F. nottii*, E) *F. olivaceus*.

For *F. xenicus* and *F. catenatus*, ONT read qualities ranged from Q5 (minimum cutoff) to Q14 with read lengths generally ranging from 10 bp to 100kb (Figure 3B,C). For *F. nottii and F. olivaceus*, ONT read qualities ranged from Q5 (minimum cutoff) to Q13 with read lengths ranging from 100 bp to 100kb (Figure 3D,E).

### Draft Assemblies

As a comparison with assemblies using long ONT read data, Illumina data alone were assembled using ABySS version 2.1.5. While the BUSCO scores were consistently above 50% completeness [46], the number of contigs and contig N50 lengths of the Illumina-only assemblies were not acceptable for downstream use (Table 3).

**Table 3.** Statistics for Illumina-only assemblies using ABySS (version 2.1.5) for each species. The BUSCO Eukaryota database (303 genes) was used to evaluate the completeness of each assembly [46]. BUSCO numbers reported are percentage complete (C) followed by the percentages of complete single-copy (CS), complete duplicated (CD), fragmented (F), missing (M) out of 303 genes.

| Species | Bases in the Illumina-only assembly | N contigs | Avg length | Largest contig | N50 | Illumina-only BUSCO C<br>CS/CD/F/M |
| --- | --- | --- | --- | --- | --- | --- |
| *F. xenicus* | 1,283,257,056 | 5,195,861 | 246.98 | 71,596 | 2,571; n = 107,350 | 57.1%<br>56.4/0.7/33.3/9.6 |
| *F. catenatus* | 1,205,429,912 | 3,989,534 | 302.15 | 70,870 | 3,629; n = 80,839 | 53.8%<br>52.8/1.0/36.0/10.2 |
| *F. nottii* | 1,167,835,004 | 3,875,693 | 301.32 | 92,540 | 3,740; n = 72810 | 62.7%<br>61.7/1.0/27.4/9.9 |
| *F. olivaceus* | 1,252,948,998 | 4,509,089 | 277.87 | 70,765 | 3,670; n = 77136 | 65.7%<br>64.0/1.7/25.1/9.2 |

The ONT-only assemblies using the fuzzy de Bruijn graph assembler, wtdbg2 (version 2.3; [47]) had high contig N50 but low complete matches with the BUSCO Eukaryota database (Table 4). The assembler wtdbg2 took an average of 6.1 hours per assembly and required 59 GB RAM. The polishing tool pilon required an average of 65.99 hours and used 1.61 TB RAM. Following polishing with Illumina data using the pilon software tool version 1.23 [33], the BUSCO Eukaryota completeness scores increased to consistently greater than 90% (Table 4). A full table of BUSCO metrics can be found in Supplemental Table 1. Assemblies were deposited in the Open Science Framework (OSF) repository, https://doi.org/10.17605/osf.io/zjv86 and in zenodo record, https://doi.org/10.5281/zenodo.3251033.

**Table 4.** ONT PromethION assemblies using the wtdbg2 version 2.3 assembler [47] followed by polishing with pilon version 1.23 [33]. Of interest is the dramatic improvement of the complete BUSCO metric after polishing with pilon. BUSCO numbers reported are percentage complete (C) followed by the percentages of complete single-copy (CS), complete duplicated (CD), fragmented (F), missing (M) out of the 303 genes in the BUSCO Eukaryota database [46].

| Species | Contigs | Contig N50 | Assembly size (bases) | Complete BUSCO after wtdbg2 ONT-only<br>C<br>CS/CD/F/M | Complete BUSCO after pilon polishing with Illumina<br>C<br>CS/CD/F/M |
| --- | --- | --- | --- | --- | --- |
| *F. xenicus* | 5,621 | 888,041;<br>n = 325 | 1,075,031,690 | 10.2%<br>10.2/0.0/11.6/78.2 | 90.5%<br>87.5/3.0/3.0/6.5 |
| *F. nottii* | 2,242 | 2,701,963;<br>n = 95 | 1,081,276,623 | 28.4%<br>11.2/0.0/22.1/66.7 | 94.4%<br>92.1/2.3/1.0/4.6 |
| *F. catenatus* | 5,854 | 436,102;<br>n = 780 | 1,163,592,740 | 11.2%<br>28.4/0.0/24.4/47.2 | 90.4%<br>88.4/2.0/2.6/7.0 |
| *F. olivaceus* | 2,622 | 2,669,230;<br>n = 105 | 1,198,526,423 | 23.4%<br>23.4/0.0/25.7/50.9 | 92.1%<br>89.8/2.3/1.3/6.6 |

## Discussion

In this study, we collected 30-45x coverage of ONT data in combination with 50-160x coverage of Illumina PE150 sequencing data and generated draft genome assemblies for four species of *Fundulus* killifish. For these assemblies, the combination of ONT and Illumina data allowed us to generate highly contiguous assemblies with acceptable BUSCO results. The assemblies generated by ONT data alone were not acceptable for use because of the low BUSCO results, due to the high rate of ONT sequence errors. Polishing the ONT assemblies with the Illumina data did not improve contiguity of the assemblies, but served to correct errors, shown by the large boost in BUSCO scores relative to the ONT assemblies alone.

The qualities of the ONT data appeared to make a difference in the contig N50 metrics of the assemblies. *F. nottii* and *F. olivaceus* both had contig N50 >2Mb, while assemblies from *F. xenicus* and *F. catenatus* had contig N50 <1Mb. *F. xenicus* and *F. catenatus* had shorter average read lengths and reads N50, on average, compared to *F. nottii* and *F. olivaceus. F. nottii*, which had the lowest yield, had higher average read lengths and reads N50 compared to the other species. *F. olivaceus*, which had the highest yield, also had a high reads N50 and average read length. Therefore, when generating ONT data for draft genome assemblies, it might matter more to have a lower yield of longer reads than a higher yield of shorter reads.

While the ONT data collected were sufficient for genome assembly of these organisms, it is worth noting that our yields were lower than those advertised on the ONT website (∼100 Gb from a single PromethION flow cell) (https://nanoporetech.com/products/promethion, accessed 06/12/2019), and read length N50 was shorter than expected based on DNA gel analysis. We observed that our samples were consistently not using pores as efficiently on the PromethION compared with other runs with samples isolated from human or mammalian samples. The reasons for this are not fully understood, but this could be due to brittle property of our hmw DNA.

The Vertebrate Genome Project (<u>VGP 2018</u>) lists standards for *de novo* long-range genome assembly that include PacBio long reads, 10x linked Illumina reads, Hi-C chromatin mapping and Bionano Genomics optical maps. These four types of data each have a high cost of generation as well as associated analysis time and computational costs. While chromatin capture and Hi-C methods produce chromosome-level assemblies of very high quality [48–51], this can significantly increase the cost of the genome sequencing project. Here, we report the pairing of high-quality short Illumina reads with error-prone long reads generated from the ONT PromethION platform to generate a draft assembly at a minimum cost. The qualities of these assemblies are not as high as compared to the standards recommended by VGP (2018) with the 3.4.2.QV40 phased metric, which requires the assembly to be haplotype phased with a minimum contig N50 of 1 million bp (1Mb), scaffold N50 of 10Mb, 90% of the genome assembled into chromosomes and a base quality error of Q40, (<u>VGP</u> 2019). However, for many research purposes these assemblies are sufficient, considering the low cost and that we have a high-quality reference genome assembly for another species within the genus [40]. For *F. olivaceus* and *F. nottii*, draft assemblies using wtdbg2 [47] and pilon polishing with Illumina data [33] had contig N50 >1 Mb, which meets the minimum requirements for assemblies in downstream synteny analyses [11].

New software tools and methods for base-calling, assembling and analyzing noisy ONT long reads are being developed at a fast rate [52,53]. Because of this fast pace of software tool development for ONT data, standard operating procedures are not available. While we use these assemblies for their intended purpose of downstream comparative evolutionary analyses, the raw data are shared here with the intent that others may use them for tool development and as new workflow pipelines, algorithms, tools, and best practices emerge.

## Conclusions

Sequencing data from the ONT PromethION and Illumina platforms combined can contribute to assemblies of eukaryotic vertebrate genomes (>1 Gb). These sequencing data from wild-caught individuals of *Fundulus* killifish species are available for use with tool development and workflow pipelines. Ongoing work from our group is comparing genomic content between *Fundulus* species to address questions about evolutionary mechanisms of osmoregulatory divergence.

## Data re-use potential

We encourage use and re-use of these data for external analyses. This collection of whole genome sequencing data from the PromethION and Illumina platforms originates from wild-caught individuals of closely-related *Fundulus* killifish species, obtained for the purpose of downstream evolutionary genomic comparative analyses. These data add to the growing set of public data available from ONT PromethION sequencing platform [25,54] which can be used for developing base-calling and assembly algorithms with this type of data.

## Availability of supporting data and materials

Raw data are available in the ENA under study PRJEB29136. Draft assembly data products and quality assessment reports are available in the OSF repository: https://doi.org/10.17605/osf.io/zjv86 and zenodo: https://doi.org/10.5281/zenodo.3251033. Scripts used for this analysis workflow are available at: https://github.com/johnsolk/ONT_Illumina_genome_assembly

## List of abbreviations

BUSCO: Benchmarking Universal Single-Copy Orthologs
ENA: European Nucleotide Archive
hmw DNA: high molecular weight DNA
ONT: Oxford Nanopore Technologies
OSF: Open Science Framework
PE: paired end
VGP: Vertebrate Genome Project

## Declarations

### Ethical Approval

UC Davis IACUC protocol #17221

### Consent for publication

Not applicable.

## Author’s Contributions

Sample extractions and library preparations were done by LKJ, RS, TG, JR. Project advising by CTB and AW. Manuscript writing and editing by LKJ, RS, TG, JR, LF, CTB, and AW.

## Acknowledgements

We thank Dr. David Duvernell at Missouri University of Science & Technology and Dr. Jacob Schaefer at the University of Southern Mississippi for generously collecting and sending fish. A special thank you goes to Dr. Charlie Johnson and Dr. Richard Metz at Texas A&M University Agrilife Research Sequencing Facility for contributing Illumina NovaSeq data from *Fundulus olivaceus*. Thanks to the instructors and participants at PoreCamp USA (June 2017) for their helpful advice.

